# Targeting C5aR1 signaling reduced neutrophil extracellular traps and ameliorates COVID-19 pathology

**DOI:** 10.1101/2022.07.03.498624

**Authors:** Bruna M. Silva, Flavio P. Veras, Giovanni F. Gomes, Seppe Cambier, Gabriel V. L. Silva, Andreza U. Quadros, Diego B. Caetité, Daniele C. Nascimento, Camilla M. Silva, Juliana C. Silva, Samara Damasceno, Ayda H. Schneider, Fabio Beretta, Sabrina S. Batah, Icaro M. S. Castro, Isadora M. Paiva, Tamara Rodrigues, Ana Salina, Ronaldo Martins, Guilherme C.M. Cebinelli, Naira L. Bibo, Daniel M. Jorge, Helder I. Nakaya, Dario S. Zamboni, Luiz O. Leiria, Alexandre T. Fabro, José C. Alves-Filho, Eurico Arruda, Paulo Louzada-Junior, Renê D. Oliveira, Larissa D. Cunha, Pierre Van Mol, Lore Vanderbeke, Simon Feys, Els Wauters, Laura Brandolini, Fernando Q. Cunha, Jörg Köhl, Marcello Allegretti, Diether Lambrechts, Joost Wauters, Paul Proost, Thiago M. Cunha

**Affiliations:** Center for Research in Inflammatory Diseases (CRID), Department of Pharmacology, Ribeirão Preto Medical School, University of São Paulo, Ribeirão Preto, São Paulo, Brazil; Graduate Program in Basic and Apply Immunology, Ribeirão Preto Medical School, University of São Paulo, Ribeirão Preto, São Paulo, Brazil; Laboratory of Molecular Immunology, Department of Microbiology, Immunology and Transplantation, Rega Institute, KU Leuven, Leuven, Belgium; Department of Pathology and Legal Medicine, Ribeirão Preto Medical School, University of São Paulo, Ribeirão Preto, São Paulo, Brazil; Hospital Israelita Albert Einstein, São Paulo, Brazil; Department of Cell and Molecular Biology, Ribeirão Preto Medical School, University of São Paulo, Ribeirão Preto, São Paulo, Brazil; Virology Research Center, Ribeirão Preto Medical School, University of São Paulo, Ribeirão Preto, São Paulo, Brazil; Divisions of Clinical Immunology, Emergency, Infectious Diseases and Intensive Care Unit, Ribeirão Preto Medical School, University of São Paulo, Ribeirão Preto, São Paulo, Brazil; Laboratory of Respiratory Diseases and Thoracic Surgery (BREATHE), Department of Chronic Diseases and Metabolism, KU Leuven, Leuven, Belgium; Laboratory for Clinical Infectious and Inflammatory Disorders, Department of Microbiology, Immunology and Transplantation, KU Leuven, Leuven, Belgium; R&D Department, Dompé Farmaceutici s.p.a., via Campo di Pile, 67100 L’Aquila, Italy; Pain Research Center, Department of Anesthesiology, University of Cincinnati College of Medicine, Cincinnati, Ohio, 45267, USA; Institute for Systemic Inflammation Research, University of Lübeck, Ratzebuger Allee 160 23562 Lübeck, Germany; Laboratory of Translational Genetics, Department of Human Genetics, VIB-KU Leuven, Leuven, Belgium; Medical Intensive Care Unit, University Hospitals Leuven, 3000 Leuven, Belgium

**Keywords:** COVID-19, C5aR1, C5a, SARS-CoV-2, Myeloid cells, Neutrophils, NETs

## Abstract

Patients with severe COVID-19 develop acute respiratory distress syndrome (ARDS) that may progress to cytokine storm syndrome, organ dysfunction, and death. Considering that complement component 5a (C5a), through its cellular receptor C5aR1, has potent proinflammatory actions, and plays immunopathological roles in inflammatory diseases, we investigated whether C5a/C5aR1 pathway could be involved in COVID-19 pathophysiology. C5a/C5aR1 signaling increased locally in the lung, especially in neutrophils of critically ill COVID-19 patients compared to patients with influenza infection, as well as in the lung tissue of K18-hACE2 Tg mice (Tg mice) infected with SARS-CoV-2. Genetic and pharmacological inhibition of C5aR1 signaling ameliorated lung immunopathology in Tg-infected mice. Mechanistically, we found that C5aR1 signaling drives neutrophil extracellular trap (NET)s-dependent immunopathology. These data confirm the immunopathological role of C5a/C5aR1 signaling in COVID-19 and indicate that antagonist of C5aR1 could be useful for COVID-19 treatment.

## Introduction

COVID-19 is the major acute global public health issue in this century. Patients with severe COVID-19 develop acute respiratory distress syndrome (ARDS) that may progress to organ dysfunction, and death (Hu et al., 2021; Berlin et al., 2020). The disease itself is a consequence of infection with the SARS-CoV-2 virus, which triggers an inflammatory response by the host organism, potentially resulting in a maladaptive inflammatory response and progression to severe disease (Valle et al., 2020; Paludan and Mogensen, 2022). As in many other human viral diseases, pathology is thus mainly a consequence of the host’s response to the virus rather than of the virus itself. Reducing viral loads after the dysfunctional immune response developed may be considered but could be a less favorable therapeutic option compared to appropriate control of inflammation. Combining antiviral with immune control, including the development of specific anti-inflammatory agents to block virus-triggered inflammatory responses, might be a strategic option to treat short-living virus-caused pathology, especially in COVID-19. This hypothesis has been confirmed by the demonstration that drugs targeting the inflammatory response are, at least in part, effective to control COVID-19 severity (van de Veerdonk et al., 2022; Chen et al., 2022; Karakike et al., 2021; Durán-Méndez et al., 2021; Shankar-Hari et al., 2021; P et al., 2021). Nevertheless, these therapies need to be used with caution since they may also affect the host immune response against the virus and against secondary/opportunistic infections. Noteworthy, the development of novel agents to treat COVID-19 targeting the inflammatory/immune response should be focused on a mediator/process that is important for immune pathology but dispensable for infection control (Risitano et al., 2020). One possible candidate might be complement C5a/C5aR1 signaling (Risitano et al., 2020).

C5a is one of the most important components of the complement cascade and possesses several pro-inflammatory actions (Risitano et al., 2020; Gerard and Gerard, 1994). C5a is a common component of the activation of all complement pathways and acts mainly via the G protein-coupled receptor (GPCR) C5a Receptor type 1 (C5aR1), also called CD88 (Gerard and Gerard, 1994). C5aR1 was initially identified in neutrophils, monocytes/macrophages, and mast cells (Gerard and Gerard, 1994; Soruri et al., 2003). The C5aR1 signaling has been implicated in the pathophysiology of several inflammatory diseases including virus-infection-induced diseases that cause lung pathology (Vandendriessche et al., 2021; Song et al., 2018; you Zheng et al., 2020; Sadik et al., 2018; Mulligan et al., 1996). For instance, C5a/C5aR1 inhibition alleviates lung damage in a murine model of influenza A, Middle East respiratory syndrome coronavirus (MERS-CoV), and respiratory syncytial virus (RSV) (Jiang et al., 2018; Garcia et al., 2013; Hu et al., 2017).

A growing body of evidence suggests the possible participation of the complement system, and especially of C5a/C5aR1 signaling in COVID-19 pathophysiology (Carvelli et al., 2020; Cugno et al., 2020). C5a levels increased in the blood of COVID-19 patients and correlated with disease severity (Carvelli et al., 2020). Nevertheless, no study investigated in depth the outcome of the lack or blockade of C5aR1 signaling on COVID-19, or the mechanisms behind its role. Herein, we found that C5a/C5aR1 signaling is increased in patients and in a preclinical mice model of COVID-19. Furthermore, we show that genetic and pharmacological blockage of C5aR1 signaling in myeloid cells (especially neutrophils) ameliorates COVID-19 lung immunopathology. Finally, we found that the C5aR1 signaling mediates COVID-19 immunopathology through enhancement of neutrophil extracellular traps (NETs) formation.

## Results and Discussion

### C5a/C5aR1 signaling in the lung cells of COVID-19 patients

COVID-19 is a consequence of two main factors: the virus replication that *per se* causes cellular injury and the dysregulated inflammatory/immune response that amplifies the tissue/organ dysfunction, especially in the lung. Although there is a race to identify novel antiviral drugs capable to inhibit SARS-CoV-2 replication and then reduce COVID-19 severity, drugs that target the inflammatory/immune response, at least partially, have been shown effective in ameliorating COVID-19 (Tomazini et al., 2020; Lopes et al., 2021; Salama et al., 2021; AC et al., 2021; Kyriazopoulou et al., 2021). Thinking about drugs targeting the immune system to control COVID-19, it is desirable to identify immune cells/mediators and molecular mechanisms that are not involved in the control of virus infection (and possible secondary infection) but are critical for immunopathology. Among several inflammatory mediators that may possess these characteristics, we and others consider complement factor C5a, and its receptor C5aR1, among the most interesting candidates (Risitano et al., 2020; Woodruff and Shukla, 2020). In agreement, targeting C5a/C5aR1 signaling ameliorates virus infection-induced lung diseases (Song et al., 2018; you Zheng et al., 2020; Sadik et al., 2018; Mulligan et al., 1996; Jiang et al., 2018; Garcia et al., 2013; Hu et al., 2017). Regarding COVID-19, the reported higher levels of C5a in the plasma of COVID-19 patients that may correlate with disease severity already are indicate for C5a levels as a possible biomarker (Cugno et al., 2020; Holter et al., 2020; Prendecki et al., 2020; Cugno et al., 2021). Moreover, it might indicate C5a plays a detrimental role in the pathophysiology of the disease. To test this hypothesis, we assessed bronchoalveolar lavage (BAL) fluid from critically ill COVID-19 patients requiring invasive mechanical ventilation, which we have previously reported to contain increased numbers of hyperactivated degranulating neutrophils and elevated concentrations of e pro-inflammatory cytokines/chemokines (eg. IL-1β, IL-6, G-CSF, CCL2, CCL3, CCL4, CXCL1 and CXCL8) compared to mechanically ventilated patients with influenza infection, as a non–COVID-19 viral pneumonia cohort (Cambier et al., 2022). We analyzed the levels of C5a in these cohorts of patient samples and found that C5a concentrations are higher in BAL fluid from COVID-19 patients than in BAL fluid from influenza-infected patients (Fig. 1a). In addition, the levels of C5a were also higher in the BAL fluid compared to the corresponding paired plasma samples (Fig. 1b). Together these results indicate that C5a is highly activated locally (in lungs) in COVID-19 and corresponds to a potential stronger local complement activation in COVID-19 compared to another viral lung infection.

**Figure 1.**
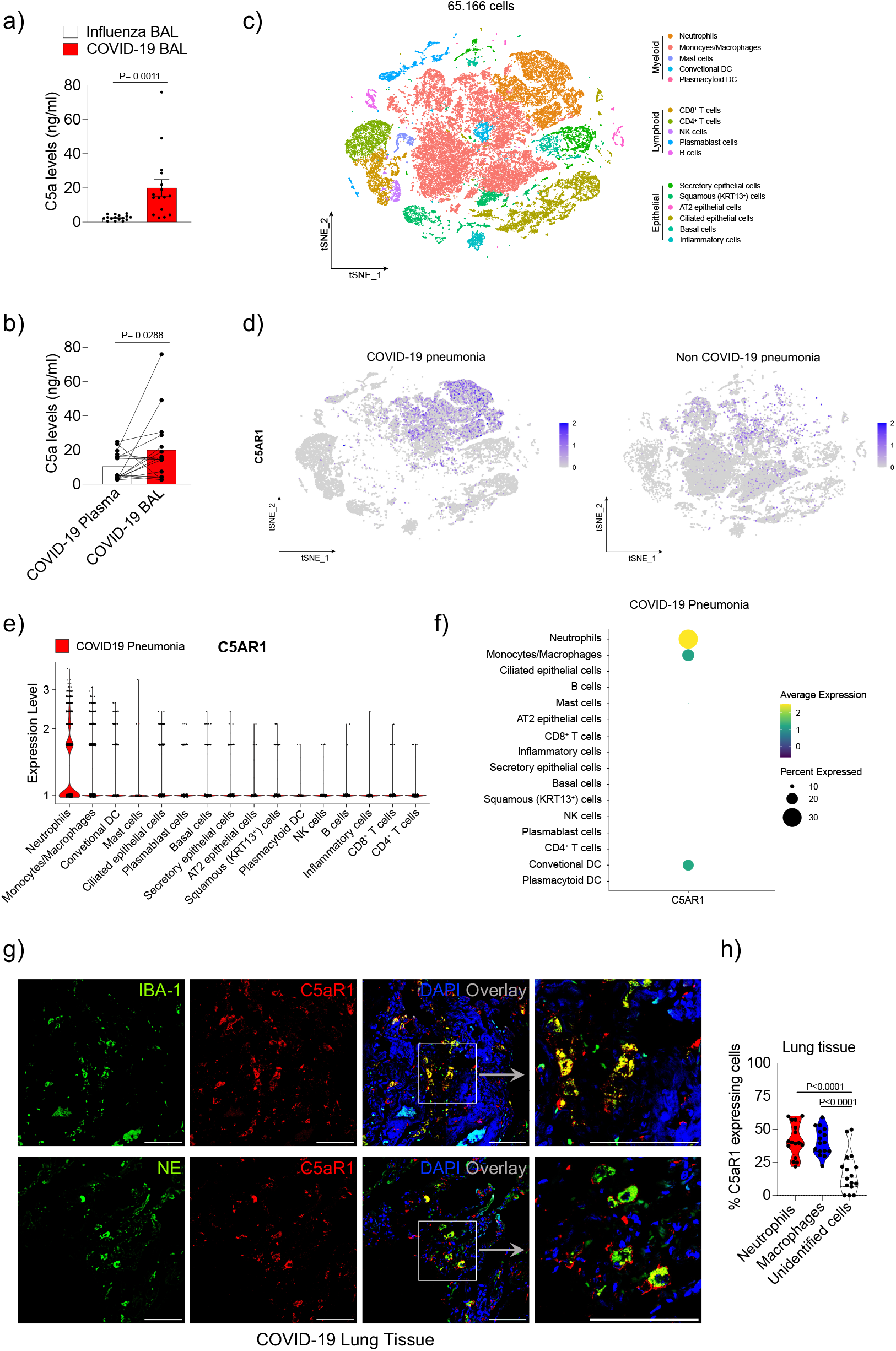
C5a levels and *C5AR1* expression in the BAL fluid and cells from COVID-19 patient. (**a**) An ELISA assay was performed to measure C5a concentrations in non-COVID-19 (n=16) and COVID-19 patients (n=16) in BAL fluid and plasma (**b**) Paired concentrations of C5a in the plasma and BAL fluid from COVID-19 patients were determined by ELISA (**c**) t-Distributed Stochastic Neighbor Embedding (t-SNE) analysis of total cells (65,166) from BAL fluid of non-COVID-19 pneumonia (n=13) and COVID-19 patients (n=22). Dot plots display the highlighted distribution of *C5AR1* each indicated cell population (**d**). (**e**) Violin plots showing the expression levels of *C5aR1* in each type of cells. The dot plot depicts the scaled and centered expression of an average cell in each cluster and therefore contains negative and positive values. The average expression reflects the mean expression of *C5AR1* in each cluster compared with all other cells (**f**). (**g**) Representative confocal images of the presence of C5aR1 in macrophages (Iba-1) and neutrophils (NE) in the lung tissue from autopsies of COVID-19 patients (n=4 cases/4 randomized field). Cells were stained for nuclei (DAPI, blue), Iba-1 or NE (green), and C5aR1 (red). Scale bar indicates 50 µm. (**h**) Percentage of cells expressing C5aR1 in the COVID-19 lung. Data are shown as the mean ± SEM. P values were determined by two-tailed unpaired (**a**) or paired (**b**) Student *t*-test followed by Wilcoxon matched-pairs signed rank test, and one-way ANOVA followed by Bonferroni’s post hoc test (**h**)

Moreover, the increased levels of C5a in the BAL fluid might indicate the activation of C5aR1 signaling. Thus, in an attempt to gain information about the possible role of C5a in the pathophysiology of COVID-19, we sought to identify the possible cells subtype in the BAL fluid of COVID-19 patients expressing *C5AR1*, its main pro-inflammatory receptor (Woodruff and Shukla, 2020; Sadik et al., 2018). To this end, we assessed our previously published database containing single-cell transcriptomes of BAL fluid cells from COVID-19 patients and from non-COVID-19 pneumonia patients and re-analyzed these data (Wauters et al., 2021). Among the different clusters of cells, we found in the BAL fluid from these patients (Fig. 1c), *C5AR1* expression is enriched in neutrophils, which are more abundant in the BAL fluid of COVID-19 patients compared to non-COVID pneumonia (Fig. 1d-f). *C5AR1* is also expressed in macrophages/monocytes in both groups of patients (Fig. 1d-f). Corroborating these data, the re-analyses of a public data set of the single-cell transcriptome of cells from BAL fluid of COVID-19 patients (Liao et al., 2020) revealed similar results (Fig. S1). Of note, both single-cell transcriptome data sets also revealed some degree of expression of *C5AR1* in epithelial cells of the BAL fluid of COVID-19 patients (Fig. 1e **and Fig. S1c**).

In order to validate the single-cell transcriptome data, lung tissue from post-mortem COVID-19 patients was used for C5aR1 immunostaining and co-staining with neutrophils (Neutrophil elastase; NE) and macrophages/monocytes (Iba-1) cellular markers. In agreement with the single-cell transcriptome data, we found C5aR1 expression mainly in cells expressing NE (neutrophils; 41.87 ± 12.77%; Fig. 1g and h) and Iba-1 (macrophage/monocytes, 40.87 ± 10.22%, Fig. 1g and h) The remaining non-identified cells were 17.49 ± 15.52% (Fig. 1h), which could be related to the epithelial cells that we found expressing *C5AR1* in the single-cell transcriptome analyses. Together these data indicate that in COVID-19 the enhanced production of C5a in the lung mainly act on neutrophils and/or macrophages/monocytes.

In an attempt to obtain further information about the possible role of C5a/C5aR1 signaling in the pathophysiology of COVID-19, we performed correlation analyses of C5a concentration with different inflammatory markers/cells that we have previously shown to be enhanced in the BAL fluid of COVID-19 patients (Cambier et al., 2022). Notably, C5a levels correlate with the number of hyperactivated/degranulating neutrophils (expressing CD66b, Sialyl-LewisX and the tetraspanin, CD63) (Fig. S2), and with CXCL8 but not with any other inflammatory marker (Fig. S2). In agreement, hyperactivated neutrophils in the BAL fluid of COVID-19 patients were characterized by higher expression of CXCL8 and they seem to play a critical role in COVID-19 pneumonia (Cambier et al., 2022; Vanderbeke et al., 2021; Cui et al., 2021; Diamond, 2020). Altogether, these data point towards a possible role for C5a in the hyperactivation of neutrophils in the lung tissue of COVID-19 patients. These results are also in accordance with the literature showing that: a) C5a/C5aR1 signaling directly triggers neutrophils chemotaxis and activation (eg. granule enzyme release and superoxide anion production/respiratory burst) in several pathological conditions (Denk et al., 2017; Ward, 2004; GERARD and GERARD, 1991; Gerard, 2003; Sadik et al., 2012; Miyabe et al., 2017); b) C5aR1 signaling induces neutrophils to degranulate (with increase in CD66 expression) in sepsis models (Schmidt et al., 2015); c) C5a levels in soluble fraction of sputum correlated positively with markers associated with worse cystic fibrosis lung disease, including neutrophil elastase, myeloperoxidase activity and DNA concentration (Hair et al., 2017).

### C5a/C5aR1 signaling on myeloid cells has a detrimental role in a murine model of COVID-19

In order to better understand the importance and role of C5a/C5aR1 signaling on the pathophysiology of COVID-19, we moved to a well-established preclinical mouse model used to study this disease, the K18-hACE2 Tg mice (Tg)-infected with SARS-CoV-2 (Winkler et al., 2020; Zheng et al., 2021) (Fig. 2a). As observed in BAL fluid from COVID-19 patients, the levels of C5a increased in the lung tissue of Tg mice infected with SARS-CoV-2 (Fig. 2b). We also observed that clinical signals (clinical score and weight loss) and pathology of the COVID-19 mouse model were associated with increased levels of pro-inflammatory cytokines/chemokines in the lung tissue of infected mice (Fig. S3a-c) as observed previously (Puhl et al., 2020; Winkler et al., 2020). The expression of C5aR1 in lung tissue of SARS-CoV-2 infected mice was also analyzed by immunofluorescence. Tg*^Flox/Flox^*mice (which contain an eGFP reporter for C5aR1 expression) were infected and the lung was collected at 5 days post infection (dpi). Similarly to what we observed in the lung tissue of COVID-19 patients, immunofluorescence analyses of the lung tissue of SARS-CoV-2-infected Tg*^Flox/Flox^* mice revealed that C5aR1 is mainly expressed in cells positive for NE (neutrophils, 41.2 ± 16.07%) and Iba-1 (macrophages, 48.62 ± 15.07%) (Fig. 2 c **and** d). The C5aR1 seems to be expressed by 10.17±6.08% of unidentified cells (Fig. 2d) These results indicate that during SARS-CoV-2 infection in mice, there is also a local activation of C5a/C5aR1 signaling especially in neutrophils and macrophages/monocytes.

**Figure 2.**
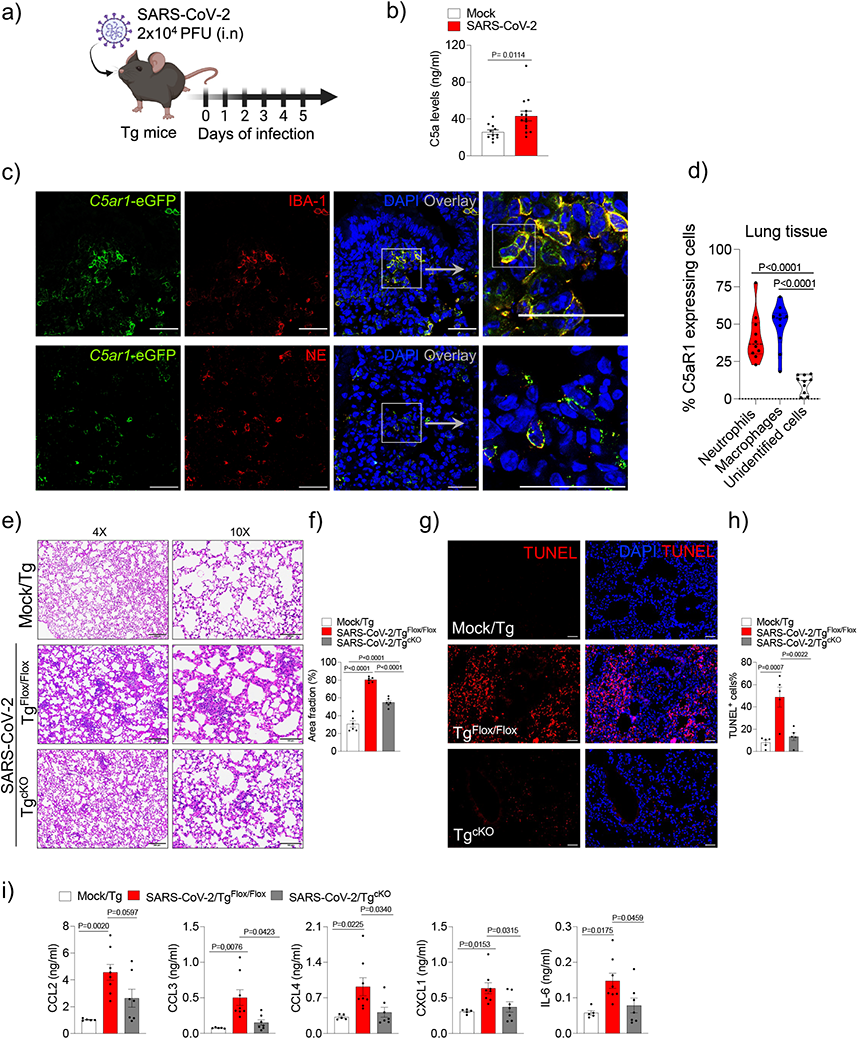
C5aR1 signaling on myeloid cells account for the lung pathology in a COVID-19 mouse model. (**a**) Tg mice were infected with SARS-CoV-2 (2 x 10^4^ PFU, i.n). (**b**) ELISA assay to measure levels of C5a in the lung tissue of infected animals (n=14) or mock control (n=11). (**c**) Representative confocal images of the presence of C5aR1 expression in the lung tissue of Tg^Flox/Flox^ mice (*C5ar1*-eGFP mice) infected with SARS-CoV-2 (5 dpi). Tissues slices were co-stained stained for nuclei (DAPI, blue), Iba-1 (macrophages, red) and NE (neutrophils, red) markers. Scale bar indicates 50 µm. (**d**) Percentage of cells expressing C5aR1 in the lung tissue of Tg^Flox/Flox^ mice infected with SARS-CoV-2 (n=4 mice/4 randomized field). (**e**) Representative H&E staining from the lung of SARS-CoV-2-infected Tg^Flox/Flox^ (n=6) or Tg^cKO^ mice (n=6). Mock was used as control (n=6). (**f**) Quantification of the lung septal area fraction. (**g**) TUNEL staining (red) for detection of apoptotic cells *in situ* from lung tissue of SARS-CoV-2-infected Tg^Flox/Flox^ (n=5) or Tg^cKO^ mice (n=6). Mock-infected Tg mice were used as a control (n=5/group). (**h**) Percentage of TUNEL positive cells in lung tissue. Scale bar indicates 50 µm. (**i**) ELISA assays were performed to detect CCL2, CCL3, CCL4, CXCL1 and IL-6 levels in the lung tissue of Tg^Flox/Flox^ (n=8) or Tg^cKO^-infected mice (n=7). Mock-infected Tg mice were used as a control (n=5). Data are shown as the mean ± S.E.M. P values were determined by (**b**) Student’ *t*-test and one-way ANOVA followed by Bonferroni’s post hoc test (**d**, **f**, **h** and **i**).

Based on the fact that the pattern of expression of C5aR1 is mainly concentrated in myeloid cells (neutrophils and macrophages/monocytes) in the lung of COVID-19 patients and Tg mouse-infected by SARS-CoV-2, we developed a colony of Tg mice that lacks C5aR1 (Tg^cKO^ mice) signaling in these immune cells and they were infected with SARS-CoV-2 (Fig. S4a). Although we did not observe any difference in the weight loss or clinical score in Tg^cKO^-infected mice compared to Tg^Flox/Flox^ mice during the course of the disease (Fig. S4b and c), the histopathological analysis of the lung revealed a reduced level of tissue damage (Fig. 2e and f). In agreement with the histopathological data, the number of TUNEL positive cells in the lung tissue of Tg^cKO^ mice was also reduced when compared with the tissue of Tg^Flox/Flox^ mice, indicating a reduction in cell death and consequently a reduction in the lung tissue damage (Fig. 2 g **and** h). The reduction in lung pathology in Tg^cKO^-infected mice was also associated with a reduction in the levels of pro-inflammatory cytokines/chemokines, especially, CCL3, CCL4, CXCL1 and IL-6 (Fig. 2i). No difference was observed in the viral load between Tg^Flox/Flox^ and Tg^cKO^-infected mice (Fig. S4c). These results indicated that C5aR1 signaling on myeloid cells is involved in the SARS-CoV-2-induced lung pathology but has no participation in the control of the virus infection.

The dissociation between clinical parameters and lung pathology in Tg^cKO^-infected mice, although discrepant, might be explained by the fact that while Tg mice infected with SARS-CoV-2 developed lung disease similar to COVID-19 patients, clinical signs that led to eventual morbidity/mortality are mainly due to the central nervous system (CNS) dysfunction (Fumagalli et al., 2022; Vidal et al., 2021). In fact, high SARS-CoV-2 burden and encephalitis have been found also in the brains of these animals (Fumagalli et al., 2022; Vidal et al., 2021; Kumari et al., 2021). This has been considered a limitation of this mouse model of COVID-19 (Fumagalli et al., 2022). Alternatively, we cannot exclude that C5aR1 signaling in cell types, beyond neutrophils/macrophages, might also play a role in the pathophysiology of COVID-19 (Posch et al., 2021; Aiello et al., 2022). For instance, C5aR1 signaling in endothelial cells was found to be a prothrombogenic effector in COVID-19 patients (Aiello et al., 2022). Thus, further study will be necessary to address the role of C5/C5aR1 signaling in cells other than myeloid cells for the pathophysiology of COVID-19.

### A pharmacological C5aR1 antagonist ameliorates COVID-19 in the mouse model

Since C5a/C5aR1 signaling seems to be involved in the immunopathology of COVID-19, we sought to test the efficacy of DF2593A, an orally-acting C5aR1 allosteric antagonist (Moriconi et al., 2014), on SARS-CoV-2-infected Tg mice to explore this candidate for the treatment of COVID-19. As a proof-of-concept experiment, we treated Tg mice with DF2593A (3 mg/kg, p.o.) 1h before SARS-CoV-2 infection and once a day up to the day of sample collection (5 dpi) (Fig. 3a). Notably, the treatment with DF2593A reduced the body weight loss and improved the clinical score (Fig. 3b) of the Tg-infected mice compared to vehicle-treated mice. This treatment also ameliorates lung pathology and reduces the number of dead cells (TUNEL+ cells) in the lung tissue of Tg-infected mice when compared to the vehicle-treated group (Fig. 3c-f), while it did not alter the viral load (Fig. S5a). Corroborating these results, *in vitro* data showed that DF2593A was also not effective to inhibit SARS-CoV-2 replication in Vero E6 cells (Fig. S5b). The reduction in lung pathology was also associated with a reduction in the levels of pro-inflammatory cytokines/chemokines, especially CCL3 and IL-6 in the lung tissue of mice treated with DF2593A (Fig. 3g). These preclinical results indicate that pharmacological inhibition of C5aR1 could be a novel approach to ameliorate COVID-19.

**Figure 3.**
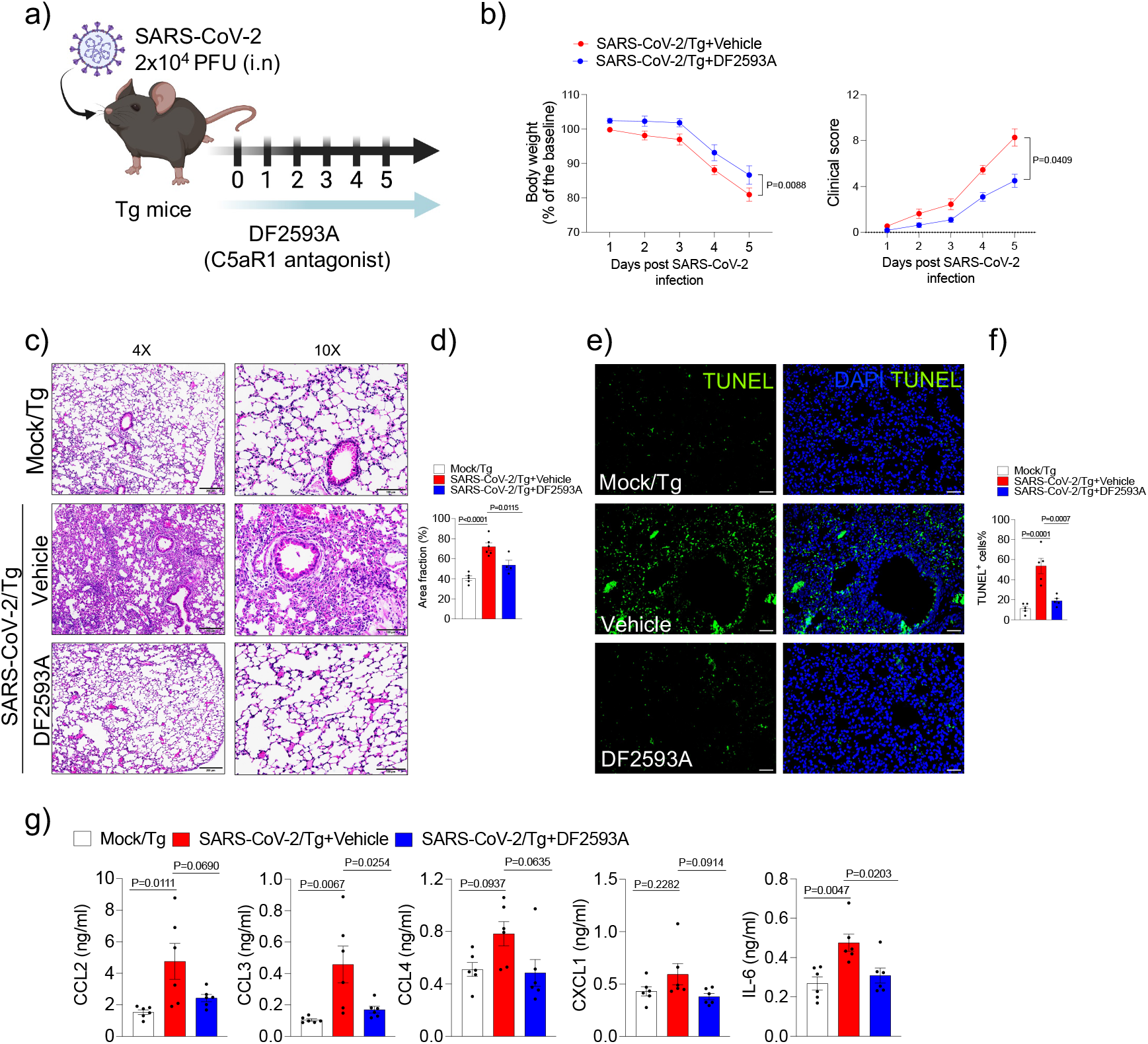
DF2593A, a selective C5aR1 antagonist, prevents lung pathology in SARS-CoV-2-infected Tg mice. (**a**) Tg mice were infected with SARS-CoV-2 (2 x 10^4^ PFU, i.n) and treated with DF2593A (3 mg/kg, p.o) until 5 dpi. (**b**) Body weight and clinical score were measured daily post-infection (n=11/group, pooled of 2 independent experiments). (**c**) Representative H&E staining from the harvested lung of the COVID-19 mouse model treated (n=4) or not (n=6) with DF2593A. Mock was used as control (n=5). (**d**) Quantification of the lung septal area fraction. (**e**) TUNEL staining (green) for detection of apoptotic cells *in situ* from lung tissue of mice (n=5/group). (**h**) Percentage of TUNEL positive cells in lung tissue. Scale bar indicates 50 µm. (**i**) ELISA assays were performed to detect CCL2, CCL3, CCL4, CXCL1 and IL-6 levels from lung (n=6/group). Mock was used as control group. Data are shown as the mean ± S.E.M. P values were determined by one-way ANOVA followed by Bonferroni’s post hoc test (**b, d**, **f** and **g**).

The higher efficacy promoted by DF2593A treatment on clinical parameters of the COVID-19 mice model compared to the phenotype observed in Tg^cKO^ mice is not immediately apparent, but it could also be explained by the fact that C5aR1 is expressed in cells other than myeloid cells, which are probably also inhibited by the C5aR1 antagonist. Additionally, since we have previously shown that DF2593A is able to cross the blood-brain barriers (Moriconi et al., 2014) it might also reduce brain inflammation which is, as we mentioned before, an important drawback of this COVID-19 mice model (Moriconi et al., 2014).

### C5a/C5aR1 signaling enhances NETs formation to aggravate COVID-19

C5a/C5aR1 signaling in myeloid cells (especially in neutrophils) is able to promote cell migration by triggering their arrest on the endothelium and/or chemotaxis (Sadik et al., 2018; Heit et al., 2002), suggesting it would be involved in the recruitment of these cells into the SARS-CoV-2 infected lungs. Thus, we further analyzed whether the lack of C5aR1 signaling in myeloid cells could impact the infiltration of these cells in the lung of SARS-CoV-2 infected Tg mice. Notably, FACS analyses revealed that the infiltration of total leukocytes (CD45+ cells), myeloid cells (CD45+CD11b+) as well as neutrophils (CD45+CD11b+Ly6G+ cells) and inflammatory monocytes (CD11b+CCR2+Ly6C+) was similar in the lung tissue of Tg^cKO^-infected mice compared to Tg^Flox/Flox^ mice (Fig. S6a-d). Like what we have found in Tg^cKO^ mice, DF2593A treatment did not reduce the infiltration of total myeloid cells, neutrophils, or inflammatory monocytes (Fig. S6f-h) in the lung tissue of Tg-infected mice. On the other hand, the total leukocytes infiltration in the lung tissue of Tg-infected mice was reduced by DF2593A treatment compared to vehicle treatment (Fig. S6e). Together, these results indicated that C5aR1 signaling on myeloid cells is not involved in the infiltration of these cells into the lung of SARS-CoV-2-infected Tg mice, which could be mediated by redundant inflammatory mediators such as chemokines (eg. CXCR2 ligands). In addition, the inhibition of C5aR1 by DF2593A in cells beyond myeloid cells, might explain the reduction of total leukocyte infiltration caused by this treatment. Thus, C5aR1 signaling on no-myeloid cells might favor, directly or indirectly, the infiltration of non-myeloid leukocytes during COVID-19 in mice and that these non-myeloid cells (eg. NK cells) might be harmful for the lung during COVID-19, as already demonstrated ^59^. This might also explain why DF2593A treated mice showed a better phenotype compared to the phenotype of Tg^cKO^-infected mice.

Our findings indicating that C5a/C5aR1 signaling in myeloid cells is involved in the lung immunopathology of COVID-19, but not in the infiltration of these cells into the lung, prompted us to hypothesize that this signaling would be involved in the local activation of these cells. Additionally, our finding that C5a levels in the BAL fluid of COVID-19 patients correlate better with hyperactivated neutrophils than with the total number of neutrophils and pro-inflammatory cytokines/chemokines (Fig. S2) also supports this hypothesis. Among the downstream mechanisms by which activated neutrophils might participate in the pathophysiology of COVID-19, the production of NETs is one of the most described (Ackermann et al., 2021). For instance, we and others have previously shown that in the lung tissue of COVID-19 patients, SARS-CoV-2 directly triggers NET-dependent lung immunopathology (Barnes et al., 2020; Zuo et al., 2020; Middleton et al., 2020; Veras et al., 2020). We also found that hyperactivated neutrophils in the BAL fluid from COVID-19 patients are enriched for NET-related genes (Vanderbeke et al., 2021). Moreover, data from our lab also showed that treatment of Tg-infected mice with NETs-degrading DNase ameliorates lung pathology (Veras et al.). Thus, we evaluated whether C5a/C5aR1 signaling would be involved in NETs formation in the lungs of SARS-CoV-2-infected Tg mice. Corroborating this hypothesis, we found that the levels of NETs in the lung tissue of Tg^cKO^-infected mice were significantly reduced compared to Tg^Flox/Flox^- infected mice (Fig. 4a and b). Furthermore, we found that the lung tissue of Tg-infected mice treated with DF2593A has low levels of NETs compared to the lung tissue from vehicle-treated mice (Fig. 4c and d). Corroborating these *in vivo* findings, we found that *in vitro* human neutrophils infected with SARS-CoV-2 produced higher levels of NETs in the presence of low concentration of recombinant C5a protein (Fig. 4e and f). These findings are in agreement with evidence that plasma from COVID-19 patients triggers NETs formation by human naive neutrophils, and this was reduced by inhibition of C5aR1 signaling (Skendros et al., 2020). Altogether, these data indicate that the induction of NETs in the lung tissue of SARS-CoV-2-infected mice might be a crucial mechanism triggered by C5a/C5aR1 signaling those accounts for the pathophysiology of COVID-19 (Fig. 4g). It is noteworthy that in Tg^cKO^ mice, C5aR1 signaling is also missing in macrophages/monocytes (McCubbrey et al., 2017). Therefore, we cannot exclude that part of the protective phenotype observed in the Tg^cKO^-infected mice would be due to inhibition of C5aR1 signaling in those cells that indirectly might also affect NETs production.

**Figure 4.**
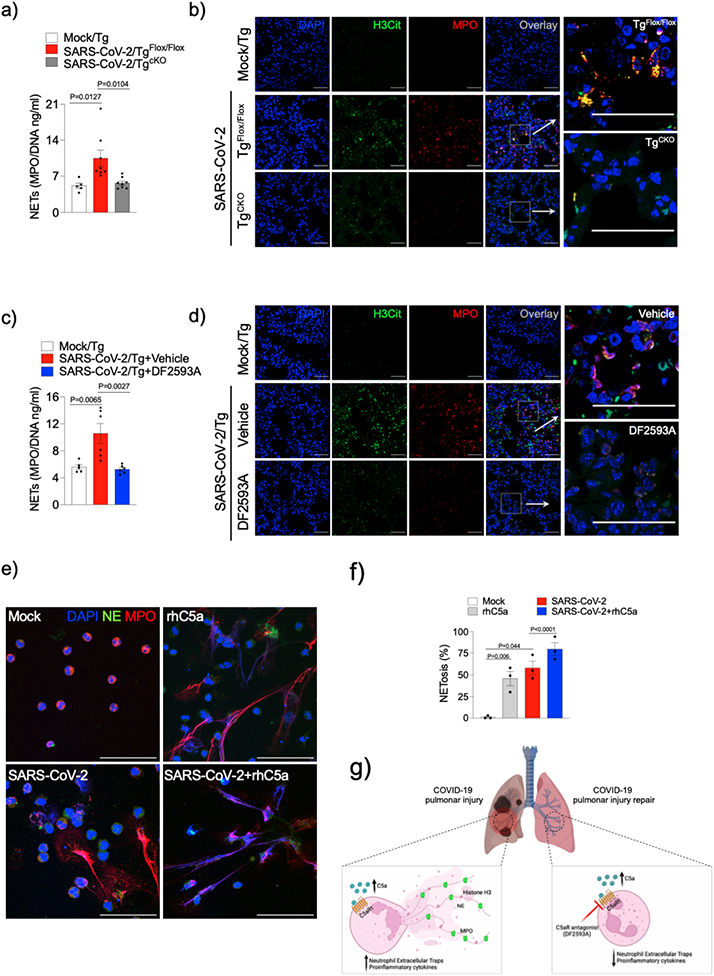
C5a enhances SARS-CoV-2-induced NET formation in neutrophils. Tg^Flox/Flox^ (n=8) and Tg^cKO^ (n=8) mice were infected with SARS-CoV-2 (2 x 10^4^ PFU, i.n). (**a**) At 5 dpi, the levels of NETs were quantified by MPO-DNA PicoGreen assay in the supernatant of the lung homogenate (**b**) Representative confocal images showing the presence of NETs in the lung tissue from Tg^Flox/Flox^ or Tg^cKO-^infected mice. A mock-infected group was performed as control (n=5). (**c**) Tg-infected mice were treated with DF2593A (3mg/kg, p.o, n=6) or vehicle (n=5/group). At 5 dpi, NETs levels were quantified by MPO-DNA PicoGreen assay in the supernatant of the lung homogenate. (**d**) Representative confocal images showing the presence of NETs in the lung tissue of Tg-infected mice treated with DF2593A or vehicle (n=5/group). Mock-infected group was performed as control (n=5). (**e**) Neutrophils were isolated from healthy controls and incubated with mock, rhC5a (3 nM) and SARS-CoV-2 (MOI = 1.0). One group of SARS-CoV-2-infected cells was pretreated with rhC5a (3 nM). (**e**) Representative images of NETs release. Cells were stained for nuclei (DAPI, blue), NE (green), and MPO (red). (**f**) Percentage of NETs quantification in these neutrophils supernatants (n=3 donors). The scale bar indicates 50 µm. (**g**) Schematic representation of the detrimental role of C5a/C5aR1 signaling on COVID-19 pathophysiology. Activation of C5aR1 signaling on myeloid cells, especially in recruited neutrophils, triggered NETs-dependent immunopathology in the lung tissue during COVID-19. Data are shown as the mean ± S.E.M. P values were determined by one-way ANOVA followed by Bonferroni’s post hoc test (**a**, **c** and **f**).

Our data reinforce the possibility to use inhibitors of C5a/C5aR1 signaling for the treatment of COVID-19. Although randomized controlled trials with modulators of this pathway for COVID-19 treatment are ongoing, no final results have been published. Nevertheless, clinical results from a small cohort indicated that inhibition of C5a reduced COVID-19 hyper-inflammation and improved lung function (Mastellos et al., 2020). Noteworthy, the hypothesis that the blockage of C5aR1 signaling would be beneficial to COVID-19 may open another important question related to secondary infections that are extremely common in COVID-19 patients and are a critical threat in the current treatments targeting the immune response (Lee et al., 2003; Hoenigl et al., 2022; Grasselli et al., 2021; Kooistra et al., 2021; Rothe et al., 2021). Although, inhibition of C5 by neutralizing antibodies has been associated with increased risk of bacterial infection due to the inhibition of the formation of the membrane attack complex, the selective targeting of C5a/C5aR1 signaling may avoid harmful anaphylatoxins-induced effects (Crew et al., 2019; Langereis et al., 2020). In fact, inhibition of C5aR1 signaling reduced the consequences of exacerbated bacterial infection such as observed in sepsis (Xu et al., 2016; Fattahi et al., 2020; Rittirsch et al., 2008; Herrmann et al., 2018). These studies gave support for the hypothesis that C5a/C5aR1 signaling is more important for immunopathology than for immune defense against infections.

Overall, our study provides direct evidence of the detrimental role C5a/C5aR1 signaling for the lung immunopathology in COVID-19. It also provides the molecular mechanism by which C5aR1 signaling, especially in neutrophils via NETs-dependent lung pathology, mediates COVID-19 pathophysiology (Fig. 4g). In conclusion, our study confirms that inhibition of C5aR1 signaling, for example by orally active allosteric inhibitors, could be an alternative against this disease.

## Material and Methods

### COVID-19 mouse model

K18-hACE2 transgenic (Tg) mice (B6.Cg-Tg(K18-ACE2)2Prlmn/J, cat. 034860) and *Lyz2*^Cre/Cre^ (B6.129P2-Lyz2tm1(cre)Ifo/J, cat. 004781) mice were purchased from Jackson Laboratory. *C5ar1*^Flox/Flox^ mice, which also express eGFP under the C5aR1 promoter, were kindly donated by Prof. Jörg Köhl ^79^. To generate Tg^cKO^ and Tg^Flox/Flox^ (littermate controls), Tg mice were bred with *Lyz2*^Cre/0^*C5ar1*^Flox/Flox^ mice. Local colonies of transgenic mice were established and maintained at the Animal Care Facility of Ribeirão Preto Medical School, University of São Paulo. Food and water were available *ad libitum* and mice kept in a controlled light-dark cycle. For COVID-19 induction, the animals received intranasal inoculation of SARS-CoV-2 (2 x 10^4^ PFU) which presents disease signs and lung pathology consistent with human disease. The manipulation of these animals was performed in Biosafety Levels 3 (BSL3) facility and approved by the ethics committee (protocol number n°66/2020).

### Human and mouse C5a levels quantification

The C5a levels were determined in the BAL fluid and plasma from sixteen critically ill adult patients with COVID-19 (<20 days in intensive care unit-ICU) and sixteen patients with influenza, as a non–COVID-19 viral pneumonia cohort. Both patient cohorts have been described previously (Cambier et al., 2022) and the C5a ELISA assay was performed using a duoset kit from R&D Systems (cat. DY2037). For mice, lung homogenate was obtained and the supernatant was collected. An ELISA was performed to detect the concentration of C5a using a kit from R&D Systems (cat. DY2150), according to the manufacturer’s instructions.

### Virus stock production

SARS-CoV-2 (Brazil/SPBR-02/2020 strain) was kindly provided by Prof. Edison Luiz Durigon (ICB-USP, Sao Paulo). The virus was propagated and titrated in Vero E6 cells in a biosafety level 3 laboratory (BSL3) at the Center for Virus Research, Ribeirao Preto Medical School (Ribeirao Preto, Brazil). Cells were cultured in DMEM medium (Corning; cat. 15-013-CVR) supplemented with 10% fetal bovine serum (FBS; GE Life Sciences; cat. SV30160.03) and antibiotic/antimycotic (Penicillin 10,000 U/ml; Streptomycin 10,000 μg/ml; Sigma-Aldrich; cat. P4333). The viral inoculum was added to Vero cells in DMEM (FBS 2%) incubated at 37 °C with 5% CO_2_ for 48 h. The cytopathogenic effect was observed under a microscope. A cell monolayer was collected, and the supernatant was stored at −70 °C. Virus titration was performed by calculating the plaque-forming units (PFU).

### Drugs and pharmacological treatment *in vivo*

For *in vivo* experiments, we used DF2593A (3 mg/kg p.o), a selective C5aR1 antagonist (Moriconi et al., 2014). For the COVID-19 mouse model, the drug was administered 1 hour before SARS-CoV-2 inoculation and daily post-infection. We assessed the daily: clinical scores (Supplementary Table 1) and body weight of each animal. At 5 days post-infection, the lungs from mock- and SARS-CoV-2 infected mice were collected. Lung lobules were collected, harvested, and homogenized in PBS with steel glass beads. The homogenate was added to TRIzol reagent (1:1; Invitrogen; cat. 15596026), for posterior viral titration via RT-qPCR, or to lysis buffer (1:1), for the ELISA assay, and stored at −70 °C. In another cohort experiment, the left lung was collected in paraformaldehyde (PFA 4%; Millipore; cat. 818715) for posterior histological assessment.

### *In vitro* SARS-CoV-2 infection

Vero E6 cells (1×10^5^) were pretreated with DF2593A at 0.01; 0.1; 1.0; 10.0 μM for 1 hour prior to SARS-CoV-2 infection at 37°C. Cells were infected at a multiplicity of infection (MOI) of 1.0 with infectious clone SARS-CoV-2 or mock with infection media to evaluate viral load by RT-PCR, 24 h post-infection. The treatment was performed in technical quadruplicate.

### SARS-CoV-2 viral load

SARS-CoV-2 detection was performed with primer-probe sets for 2019-nCoV_N1 and N2 (Integrated DNA Technologies; cat. 10006713), according to the US Centers for Disease Control (CDC) protocol by RT-PCR, using total nucleic acids extracted with Trizol reagent from cell pellet or lung tissue to determine the genome viral load. All RT-PCR assays were done using the Viia 7 Real-time PCR System (Applied Biosystems). A standard curve was generated in order to obtain the exact number of copies in the tested sample. The standard curve was performed using an amplicon containing 944 bp cloned in a plasmid (PTZ57R/T CloneJetTM Cloning Kit Thermo Fisher®), starting in the nucleotide 14 of the gene N. To quantify the number of copies, a serial dilution of the plasmid in the proportion of 1:10 was performed. Commercial primers and probes for the N1 gene and RNAse P (endogenous control) were used for the quantification (2019-nCov CDC EUA Kit, Integrated DNA Technologies), following the CDC’s instructions.

### Re-analysis of scRNA-seq data sets

We re-analyzed single-cell transcriptomic data from BAL fluid cells from patients with severe COVID-19 and their respective control groups (Wauters et al., 2021; Liao et al., 2020). The dataset was downloaded and the RDS file was imported into R environment version v4.04 and Seurat v4.1.1 (Stuart et al., 2019) by filtering genes expressed in at least 3 cells and more than 200 unique molecular identifiers (UMI) counts per cell. For the pre-processing step, outlier cells were filtered out based on three metrics (library size < 60000, number of expressed genes between 200 and 7500, and mitochondrial percentage expression < 20). The top 3,000 variable genes were then identified using the ‘vst’ method using the FindVariableFeatures function. Percent of mitochondrial genes was regressed out in the scaling step, and Principal Component Analysis (PCA) was performed using the top 3,000 variable genes with 40 dimensions. Additionally, a clustering analysis was performed on the first 7 principal components using a resolution of 2 followed by t-Distributed Stochastic Neighbor Embedding (tSNE), a dimensionality reduction technique for data visualization. Then, differential gene expression analysis was performed using FindAllMarkers function with default parameters to obtain a list of significant gene markers for each cluster of cells. To account for the frequency of cells expressing C5AR1, we filtered cells with raw counts of C5AR1>0. The dataset generated by authors is publicly available at: 1 <u>the EGA European Genome-Phenome Archive</u> database (EGAS00001004717 accessible at: https://ega-archive.org/studies/EGAS00001004717) or at http://covid19.lambrechtslab.org/.; 2) at https://covid19-balf.cells.ucsc.edu/.

### Human lung samples from autopsies

We performed adapted minimally invasive autopsies from 4 COVID-19 fatal cases (Duarte-Neto et al., 2020). (Supplementary table 2). Briefly, a mini-thoracotomy (3 cm) was done under the main area of lung injury identified by prior ultrasound-guided. The lung parenchyma was clamped by Collins Forceps, cut, and fixed in 10% buffered formalin (Sigma-Aldrich; cat. 252549). Minimally invasive autopsies were approved by the Ribeirão Preto Medical SchoolEthical Committee (protocol no. 4.089.567).

### H&E staining and lung pathology

Lung slices (5 micrometers) were fixed with PFA 4%, paraffin-embedding, and submitted to Hematoxylin and Eosin staining. A total of 10 photomicrographs at 50x or 200x magnification per animal were randomly obtained using the microscope Leica DMI6000B (Leica microsystems). The total septal area and the total lung area were quantified using the Pro Plus 7 software (Media Cybernetics, Inc., MD, USA). The ratio between total septal area and the total lung area was expressed as area fraction (%). The area fraction values were used for statistical comparison between groups and for graphical representation Morphometric analysis was performed in accordance with the protocol established by the American Thoracic Society and European Thoracic Society (ATS/ERS) (Hsia et al., 2010).

### Immunostaining and confocal

Lung samples from COVID-19 autopsies or Tg^Flox/Flox^-infected mice were fixed with PFA 4%. After dehydration and paraffin embedding, 5-μm sections were prepared. The slides were deparaffinized and rehydrated by immersing the through Xylene (Labsynth; cat. 00X1001.06.BJ) and 100% Ethanol (Labsynth; cat. 00A1084.07.BJ) for 15 min with each solution. Antigen retrieval was performed with 1.0 mM Ethylene Diamine Tetra Acetic acid (EDTA; Labsynth; cat. 00E1005.06.AG) 10 mM Trizma-base (Sigma-Aldrich; cat. T1503), pH 9.0 at 95°C for 30 min. Afterward, endogenous peroxidase activity was quenched by incubation of the slides in 5% H_2_O_2_ in methanol (Millipore; cat. 106009) at RT for 20 min. After blocking with IHC Select Blocking Reagent (Millipore, cat. 20773-M) at RT for 2 h, primary antibodies were incubated overnight at 4°C: mouse monoclonal anti-C5aR1 (clone: S5/1; Millipore; cat. MABF1980; 1:50), rabbit polyclonal anti-IBA1 (FUJIFILM Wako Pure Chemical Corporation; cat. 016-20001; 1:200), rabbit polyclonal anti-Neutrophil Elastase (anti-NE; Abcam; cat. ab68672; 1:100), goat polyclonal anti-myeloperoxidase (anti-MPO, R&D Systems, cat. AF3667, 1:100) and rabbit polyclonal, anti-histone H3 (H3Cit; Abcam; cat. ab5103; 1:100). The slides were washed with TBS-T (Tris-Buffered Saline with Tween 20) and incubated with secondary antibodies alpaca anti-mouse IgG Alexa Fluor 594 (Jackson ImmunoResearch; cat. 615-585-214; 1:1,000), donkey anti-goat IgG Alexa Fluor 488 (Abcam, cat. ab150129), alpaca anti-rabbit IgG AlexaFluor 488 (Jackson ImmunoReseacher; Cat. 611-545-215; 1:1,000) and alpaca anti-rabbit IgG AlexaFluor 594 (Jackson ImmunoReseacher; Cat. 611-585-215; 1:1,000). Autofluorescence was quenched using the TrueVIEW Autofluorescence Quenching Kit (Vector Laboratories, cat. SP-8400-15). The percentage of cells expressing C5aR1 was determined by colocalization between Iba1 (macrophage) or NE (neutrophil) with C5aR1 expression. Four randomized fields from four COVID-19 fatal cases or Tg^Flox/Flox^-infected mice were analyzed.

For NETs detection *in vitro*, neutrophils were plated in 24-well plates containing glass coverslips covered with 0.01% poly-L-lysine solution (Sigma-Aldrich; cat. P8920), fixed with PFA 4% at RT for 10 min, 2% bovine serum albumin (BSA; Sigma-Aldrich; cat. A7906) and 22.52 mg/ml glycine (Sigma-Aldrich; cat. G8898) in PBST (Phosphate Buffer Saline + 0.1% Tween 20) at RT for 2 h. The coverslips were stained with the following antibodies: rabbit polyclonal anti-Neutrophil Elastase (anti-NE; Abcam; cat. ab68672; 1:500), mouse monoclonal anti-MPO (2C7; Abcam; cat. ab25989, 1:800). After this, samples were washed in PBS and incubated with secondary antibodies: alpaca anti-mouse IgG AlexaFluor 488 (Jackson ImmunoReseacher; Cat. 615-545-214; 1:1,000) and alpaca anti-rabbit IgG AlexaFluor 594 (Jackson ImmunoReseacher; Cat. 611-585-215; 1:1,000). Slides were then mounted using ProLong™ Diamond Antifade Mountant with DAPI (Molecular Probes™, Thermo Fischer Scientific, Cat.P36962). Images were acquired by Axio Observer combined with LSM 780 confocal microscope (Carl Zeiss) at 630X magnification at the same setup of zoomed, laser rate and scanned with 4 fields/image (tile scan function). NETs were quantified, by the total number of cells per field versus the number of NETosis (cells with loss of nucleus segmentation, cells in the process of releasing chromatin in networks). Images acquired and analyzed using Fiji by Image J.

### Apoptosis TUNEL assay

Frozen lung tissue slices were used for the detection of apoptotic cells *in situ* with Click-iT Plus TUNEL Assay Alexa Fluor 488, according to the manufacturer’s instructions (Thermo Fisher Scientific; cat. C10617). The slides were counterstained with Vectashield Antifade Mounting Medium with DAPI. Images were acquired by microscope Leica DMI6000B (Leica microsystems) at 200x magnification at the same laser rate. Images acquired were analyzed using Fiji by Image J. The apoptosis quantification was performed as the percentage of TUNEL positive cells from DAPI staining.

### Production of NETs by isolated human neutrophils

Blood samples were collected from healthy controls by venipuncture for in vitro experiments. Neutrophils were isolated and purified by Percoll density gradient (72%, 63%, 54%, and 45%) (GE Healthcare; Cat. 17-5445-01). Isolated neutrophils were resuspended in RPMI 1640 (Corning; cat. 15 040-CVR). Neutrophil purity was >95% as determined by Rosenfeld’s Color Cytospin (Laborclin; cat. 620529). A total of 1×10^6^ isolated neutrophils were attached to coverslips coated with poly-L-lysine solution 0.1% (Sigma-Aldrich; cat. P8920) incubated for 4 h at 37°C for NET immunostaining. Neutrophils were incubated with Mock, recombinant human Complement C5a Protein (rhC5a; 3 nM; R&D; cat. 2037-C5-025/CF) or SARS-CoV-2 (MOI = 1.0). One group of cells incubated with SARS-CoV-2 was pretreated with rhC5a. The concentration of rhC5a was based on a concentration-response curve.

### NETs quantification in the lung tissue

The 96 well black plates were coated with anti-MPO antibody (Thermo Fisher Scientific; cat. PA5-16672) (1:1000) overnight at 4°C. Subsequently, the plate was washed with PBS+0.1% Tween 20 and blocked with 2% BSA for 2h at RT. The lung tissue homogenates were obtained and centrifuged at 10,000 g at 4 °C for 10 min. Then, the supernatant was collected and incubated overnight at 4°C. On the third day, MPO-bound DNA (NETs) was quantified using the Quant-iT PicoGreen kit (Invitrogen; cat. P11496) as previously described (Veras et al., 2020).

### Flow cytometry analysis

Lung tissue was harvested and digested with type 2 collagenase (1 mg/ml, Worthington; cat. LS004177) for 45 min at 37°C to acquire cell suspensions. Total lung cells (1 x 10^6^) were then stained with Fixable Viability Dye eFluor 780 (Invitrogen; cat. 65–0865-14; 1:1000) and monoclonal fluorochrome-stained antibodies specific for CD45 (BD Pharmingen; clone 30F-11; cat. 553080; 1:200), CD11b (Biolegend; clone M1/70; cat. 101212; 1:200), Ly6G (Biolegend; clone 1A8; cat. 127606;1:200), CCR2 (Biolegend; clone: SA203G11; cat. 150605;1:200), Ly6C (eBioscience; clone: HK1.4; cat. 45-5932-82; 1:200), for 30 min at 4°C. Data were acquired on FACSVerse flow cytometer (BD Biosciences) and analysis were performed using FlowJo (TreeStar) software. Gating strategies for flow cytometry analysis are schematically represented in (Fig. S7).

### Cytokines and chemokines quantification

Lung homogenate was added to the RIPA buffer in the proportion of 1:1, and then centrifuged at 10,000 g at 4 °C for 10 min. The supernatant was collected and stored at −70 °C until use. The sandwich ELISA method was performed to detect the concentration of cytokines and chemokines using kits from R&D Systems (DuoSet), according to the manufacturer’s instruction. The following targets were evaluated: CCL2, CCL3, CCL4, CXCL1, CXCL2, IFN-β, IL-6, IL-10, and TNF.

### Statistics

Statistical significance was determined by either one or two-tailed unpaired and paired Student t-test, one-way or two-way ANOVA followed by Bonferroni’s post hoc test. Spearman correlation analysis was performed by calculating a repeated measures correlation coefficient (r-value) and plotted utilizing a simple linear regression line. P<0.05 was considered statistically significant. Statistical analyses and graph plots were performed and built with GraphPad Prism 9.3.1 software.

### Data availability

The data supporting the findings are available within the paper and its supplementary information files or otherwise stated. Source data file is provided with this paper.

## Acknowledgments

The authors gratefully acknowledge the technical assistance of Ieda R. Schivo, Ana Katia dos Santos, Jenna M. Turner, Sergio R. Rosa, Diva A. Sousa, Eleni Tamburus, Marcella Daruge Grando, Soraya Jabur and Andreia Nogueira.

## Author contributions

T.M.C, B.M.S, F.P.V and G.F.G designed, performed experimental work, analyzed data, and prepared the manuscript. B.M.S, F.P.V, G.F.G, D.B.C, D.C.N and G.V.L.S performed experimental work related to FACS and analyzed data. B.M.S, F.P.V and G.F.G performed experiments related to infection and harvested tissue. S.C and F.B performed experiments with BAL samples. B.M.S performed ELISA assay. G.V.L.S, I.M.S.C, PVM, H.I.N, and D.L. performed the single-cell transcriptome analysis. A.H.S, J.C.S and C.M.S performed neutrophil isolation and NETs quantification. F.P.V and J.C.S performed immunostaining and confocal analysis. F.P.V performed TUNEL assay. S.S.B and A.T.F contributed to lung autopsy analysis and histopathological analyses. B.M.S, S.D, I.M.P and R.M performed SARS-CoV-2 viral load and viral stock. B.M.S performed *in vitro* infections. A.U.Q and J.K performed Tg^cKO^ mice generation. P.L-J, R.D.O, P.P, E.W, L.V, S.F and J.W contributed to the collection of clinical specimens and demographic and clinical characteristics analysis from COVID-19 and influenzas patients. T.R, A.S, D.S.Z, L.O.L, J.C.A-F, E.A, L.D.C, L.B, F.Q.C performed experiments and important scientific comments. M.A, L.B, E.A, F.Q.C provided critical materials and comments. T.M.C designed, directed, and supervised the study, interpreted data, and wrote the manuscript. All authors reviewed the manuscript and provided final approval for submission.

## Supplementary material

Supplementary material is available online.

## Supplementary Materials

**Figure S1.**
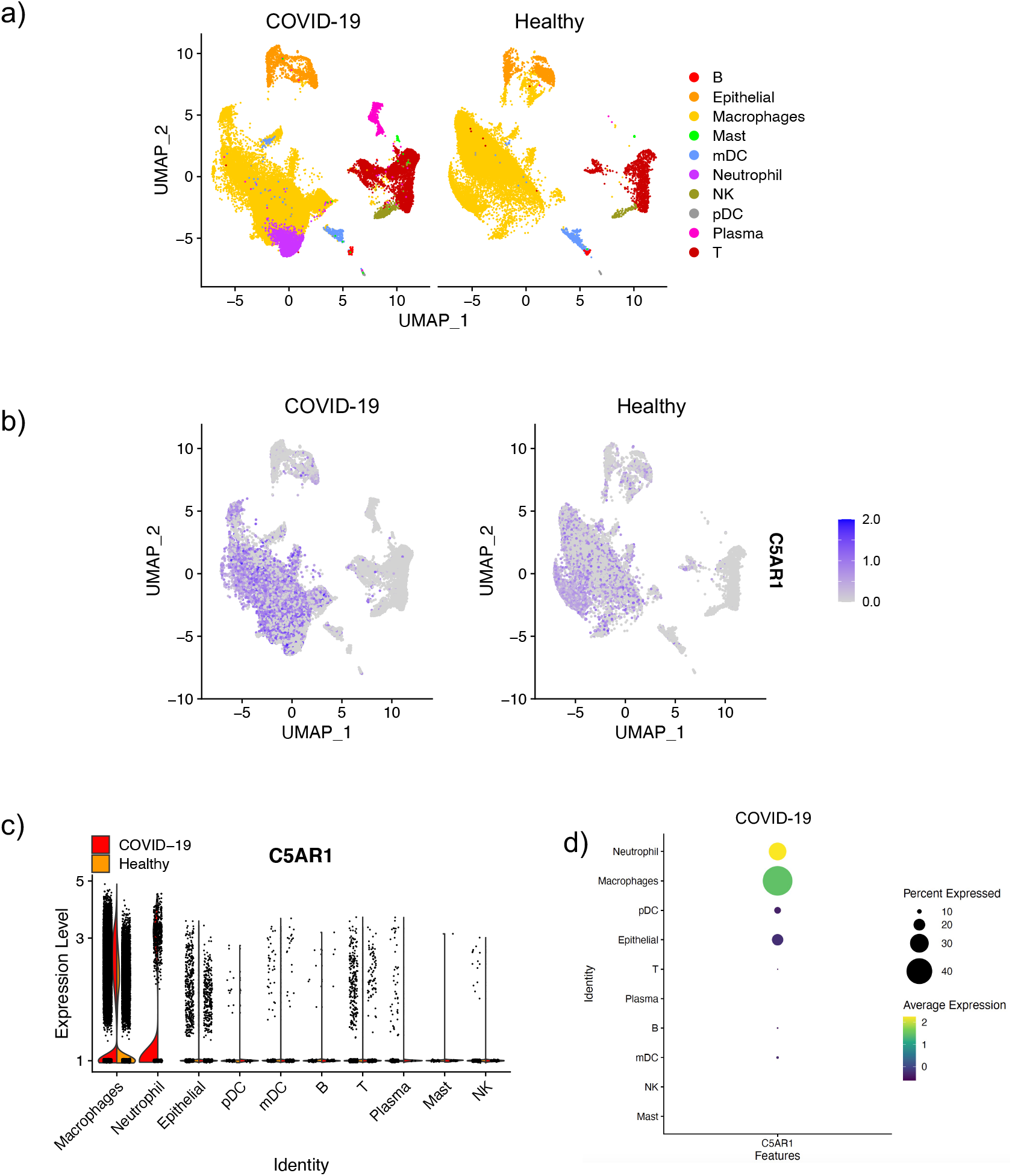
C5AR1 expression in the BAL fluid cells of COVID-19 patients. (**a**) t-SNE analysis of total cells from BAL fluid from healthy controls (n=3) and severe COVID-19 patients (n=6). (**b**) Dot plots display the highlighted distribution of *C5AR1* for each indicated cell population. (**c**) Violin plots showing the expression levels of *C5AR1* in each type of cells. (**d)** The dot plot depicts the scaled and centered expression of an average cell in each cluster and therefore contains negative and positive values.

**Figure S2.**
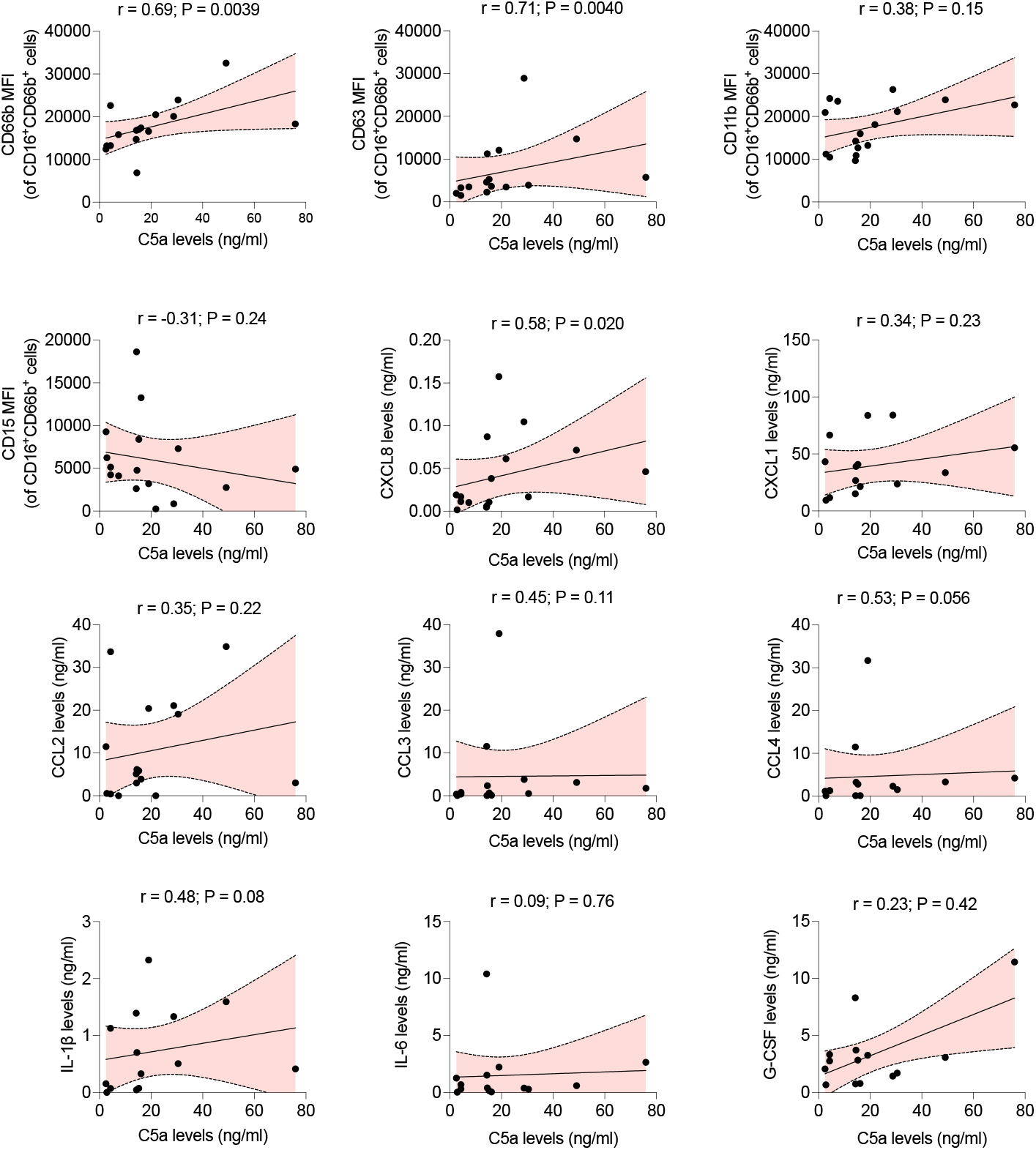
Correlation analyses between C5a and inflammatory markers in the BAL fluid from COVID-19 patients. Correlation analyses were performed with obtained levels of C5a (Fig. 1a) and the levels of different inflammatory markers that we found increased in the BAL fluid from COVID-19 patients determined in our previous study ^33^. Data depicts Spearman’s correlation coefficient (r) and P-value is depicted in each graph.

**Figure S3.**
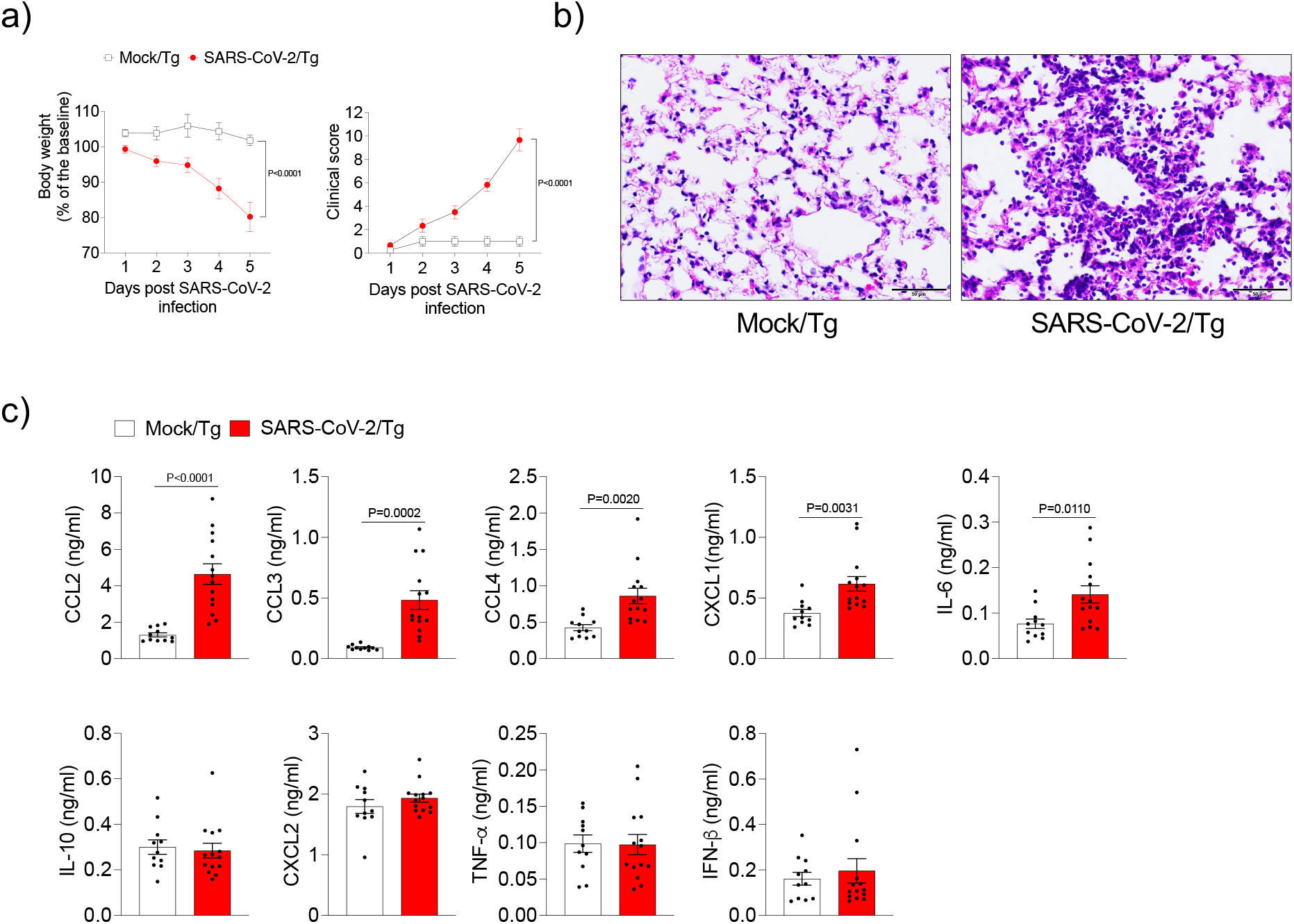
Immunopathology of COVID-19 mouse model. Tg mice were infected with SARS-CoV-2 (2 x 10^4^ PFU, i.n.) at day 0. (**a**) Body weight and clinical score were measured daily post-infection (n=4-6). (**b**) Representative image of the lung tissue of the COVID-19 mouse model harvested at 5 dpi and stained with H&E. Mock-infected group was used as control. (**c**) ELISA assays were performed to detect CCL2, CCL3, CCL4, CXCL1, IL-6, IL-10, CXCL2, TNF and IFN-β levels in lung tissue of SARS-CoV-2 infected mouse. (n=14; pooled of two independent experiments). Mock was performed as control (n=11; pooled of two independent experiments). Data are shown as the mean ± S.E.M. P values were determined by two-way ANOVA followed by Tukey’s multiple comparisons test (**a**) and unpaired Student *t*-test (**c**).

**Figure S4.**
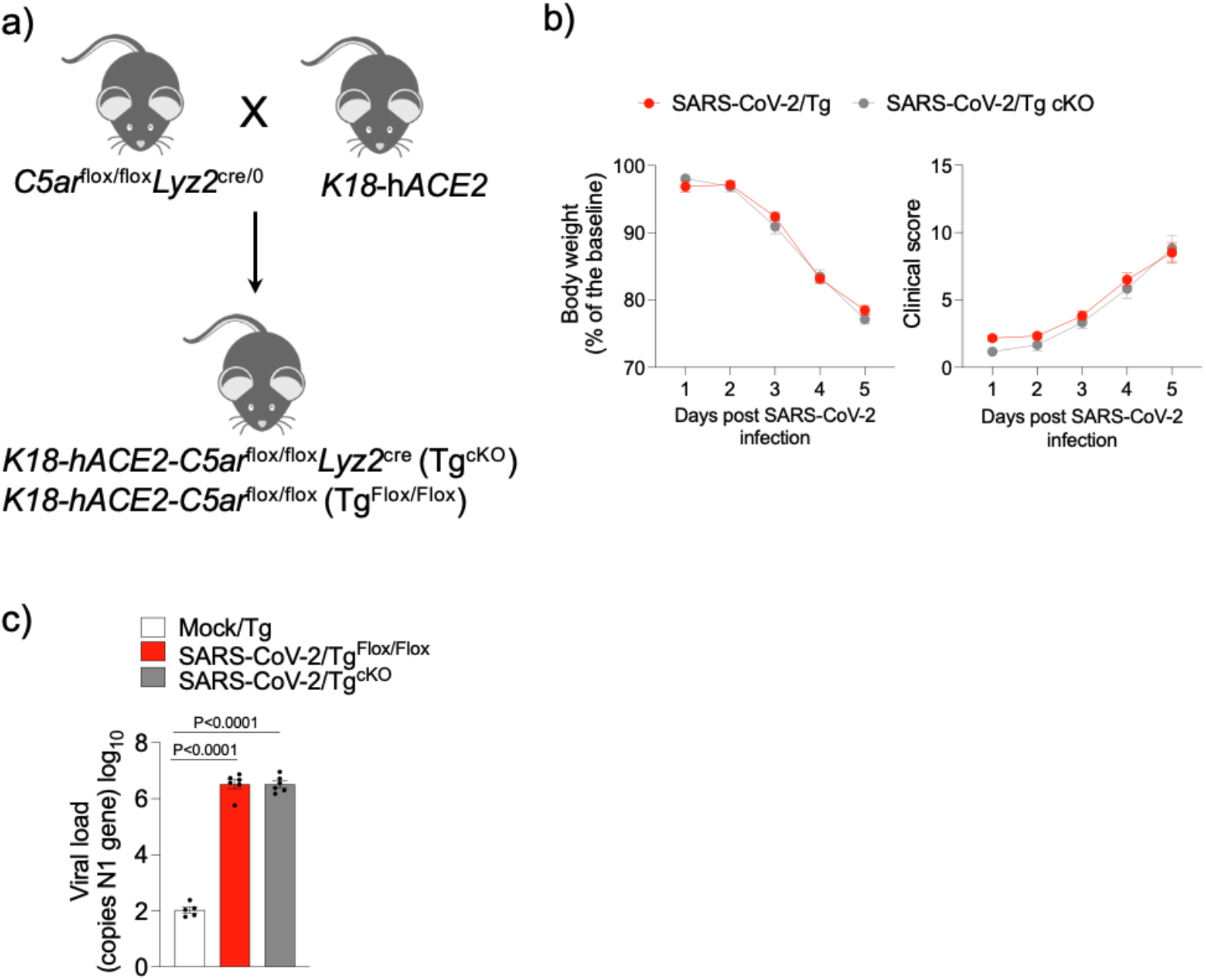
SARS-CoV-2-infected C5aR1-deficient K18-hACE2 mice. **(a)** Schematic representation of the generation of Tg^cKO^ mice and Tg ^Flox/Flox^ littermates. Body weight and clinical score were measured daily (n=6/group) in Tg^Flox/Flox^ and Tg^cKO^ mice infected with SARS-CoV-2 (2 x 10^4^ PFU, i.n.) (**b**). Viral load in the lung tissue from infected Tg^Flox/Flox^ and Tg^cKO^ mice (n=6/group) was evaluated by RT-PCR (**c**). Mock-infected group was performed as control (n=5). Data are shown as mean ± S.E.M. P-values were determined by one-way ANOVA followed by Bonferroni’s post hoc test (**c**).

**Figure S5.**
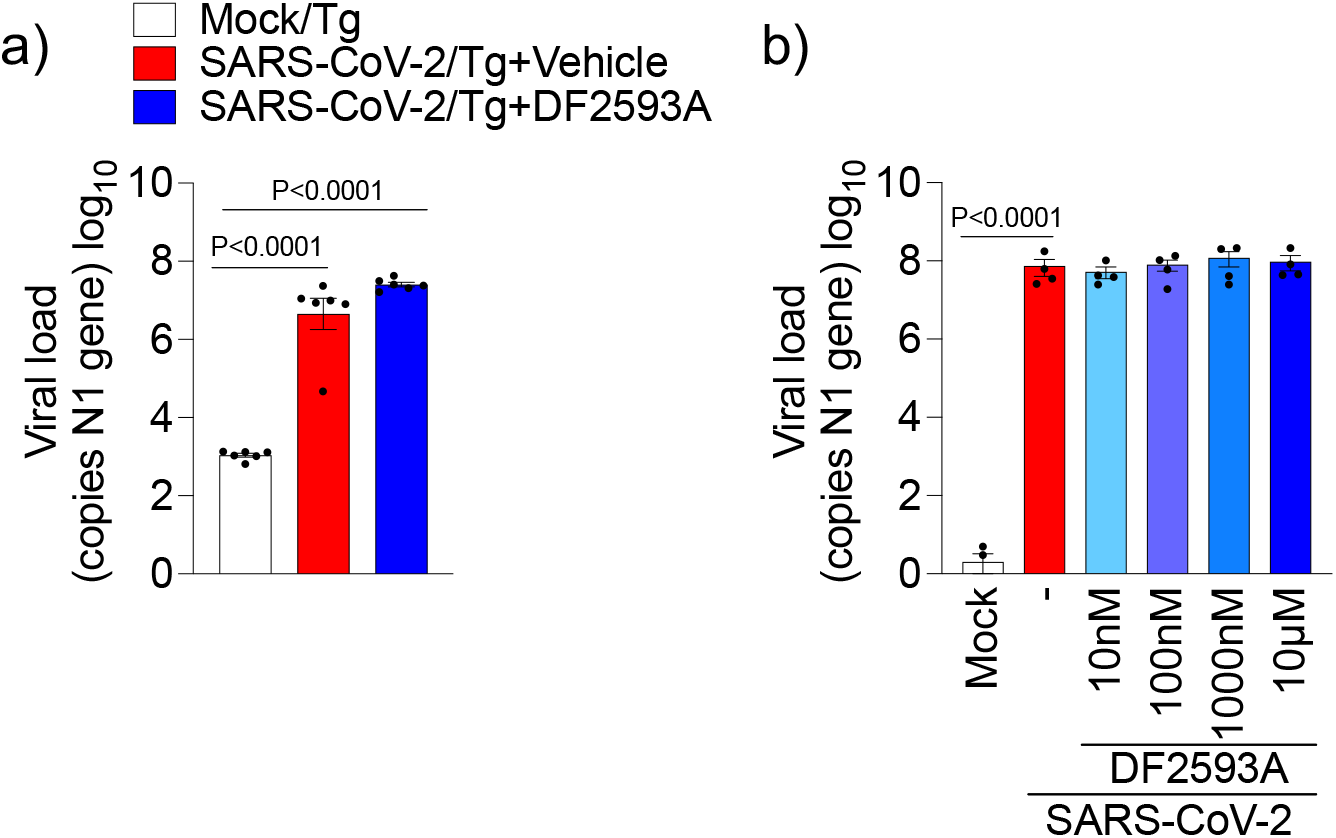
Effect of DF2593A treatment on viral load in COVID-19 mouse model. SARS-CoV-2 viral load in the lung tissue from Tg-infected mice treated with DF2593A (n=6) or vehicle (n=6). Mock-infected group was performed as control (n=6). (**b**) Vero E6 cells were pretreated with DF2593A (0,01-10 μM) followed by SARS-CoV-2 infection (MOI 1.0). Cells were lysed and the levels of SARS-CoV-2 were determined 24 h post-infection. The copy numbers of the N gene of SARS-CoV-2 was used to access the viral load by RT-PCR. Data are shown as the mean ± S.E.M. P-values were determined by one-way ANOVA followed by Bonferroni’s post hoc test (**a** and **b**).

**Figure S6.**
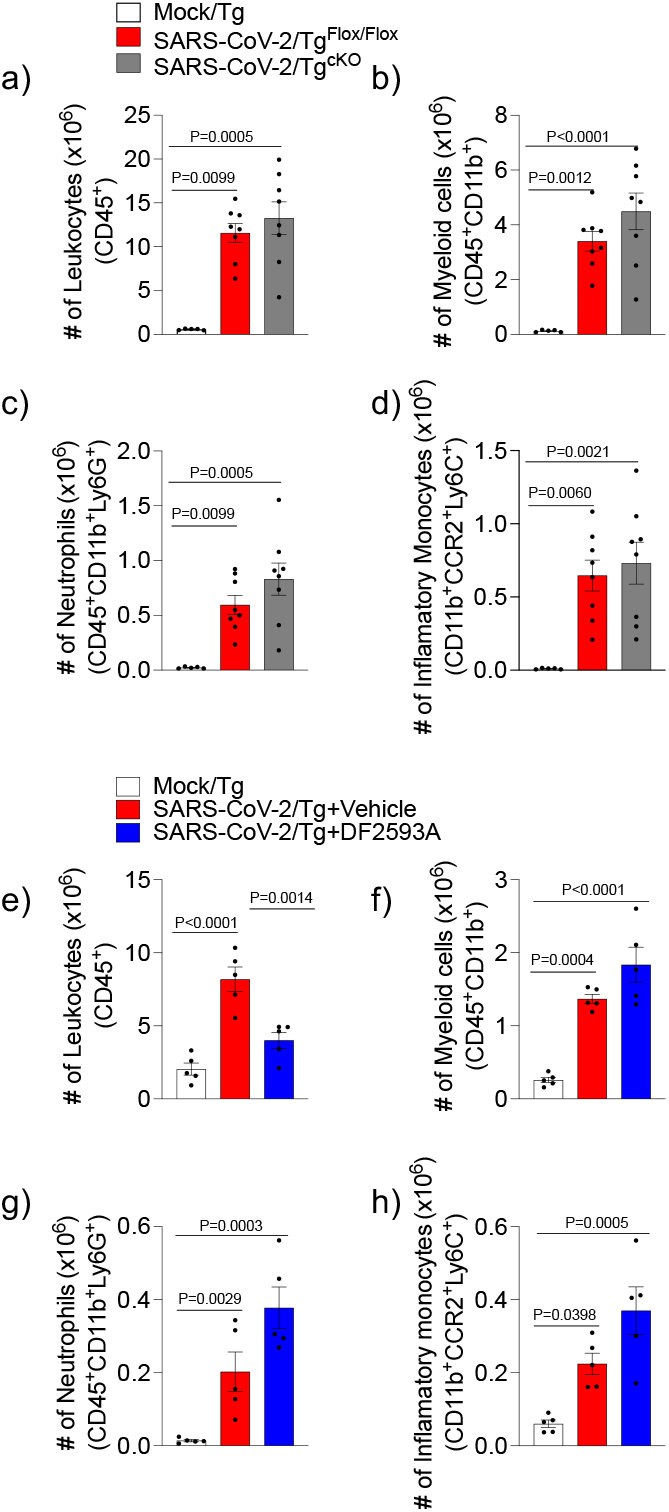
Leukocyte infiltration in the lung tissue of the COVID-19 mouse model. (**a**) Number of total leukocytes (CD45+ cells), (**b**) myeloid cells (CD45+CD11b+ cells), (**c**) granulocytes (CD45+CD11b+ Ly6G+ cells) and (**d)** inflammatory monocytes (CD45+CD11b+ CCR2+Ly6C+ cells) were determined by flow cytometry in the lung tissue from Tg^cKO^ or Tg^Flox/Flox^ infected mice harvested at 5 dpi (n= 8/per group). Mock-infected group was used as control (n=5). (**e**) Number of total leukocytes (CD45+ cells), (**f**) myeloid cells (CD45+CD11b+ cells), (**g)** granulocytes (CD45+CD11b+Ly6G+ cells) and (**h)** inflammatory monocytes (CD45+CD11b+CCR2+Ly6C+ cells) were determined by flow cytometry in the lung tissue from Tg-infected mice treated with vehicle or DF2593A (n= 5/per group) harvested at 5 dpi. Mock-infected group was used as control (n=5). Data are shown as the mean ± S.E.M. P-values were determined by one-way ANOVA followed by Bonferroni’s post hoc test.

**Figure S7.**
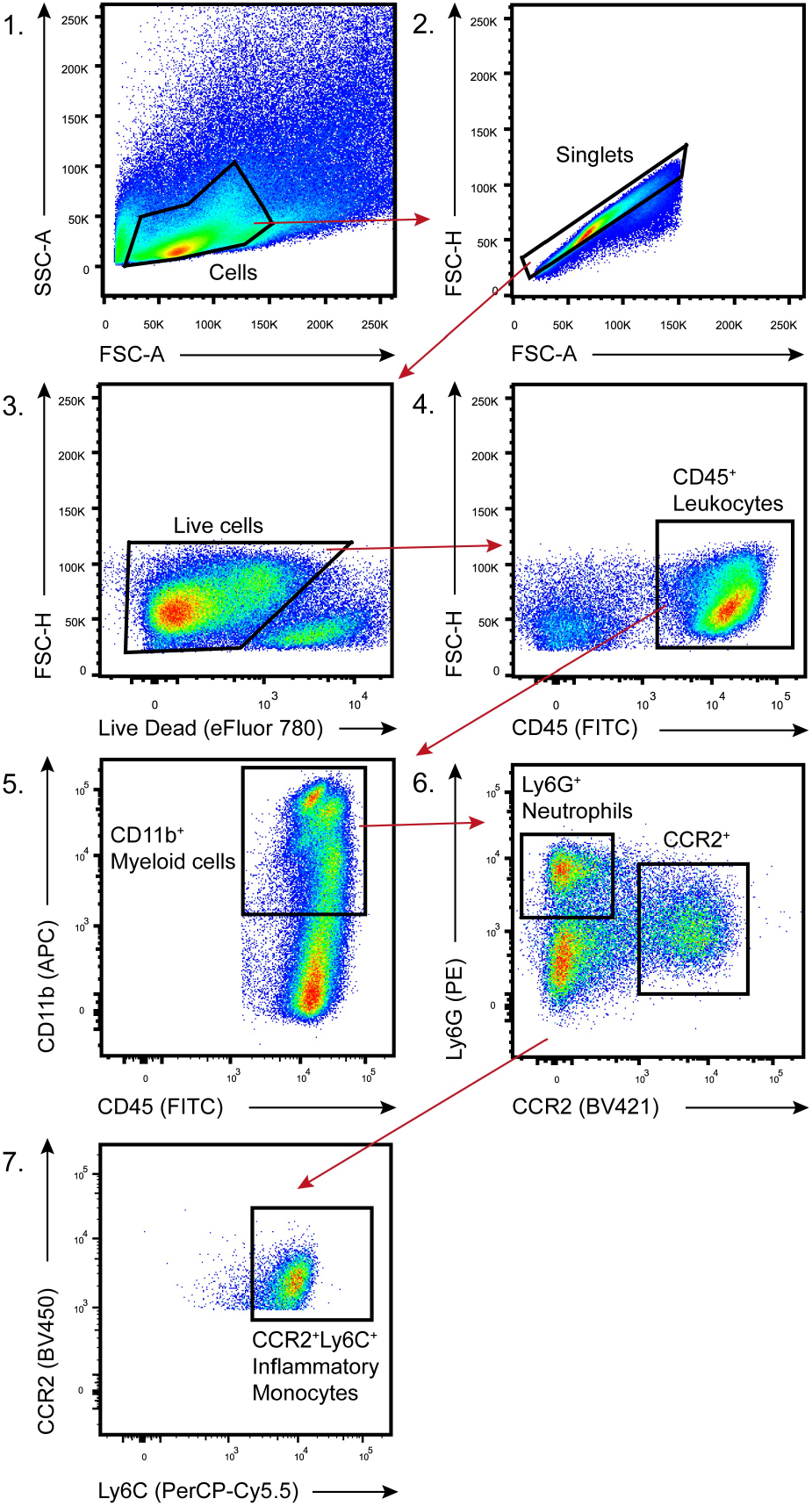
Representative gating strategies for flow cytometry analysis in the lung tissue. Gating strategies for different cell populations for all flow cytometry analyses.: **1)** Side scatter area (SSC-A) and forward scatter area (FSC-A); 2) Doublet cells were excluded by forward scatter height (FSC-H) and FSC-A gating; **3)** Viable cells were identified using fixable viability dye and FSC-H gating; **4)** the number of total leukocytes were identified as cells stained for CD45^+^ among live cells. **5)** Myeloid cells were characterized as stained for CD11b^+^ among CD45^+^ live cells. **6)** Neutrophils were identified by Ly6G^+^ inside CD11b^+^ myeloid cells. **7)** Inflammatory monocytes were characterized by Ly6C^+^CCR2^+^ among CD11b^+^ myeloid cells.

**Table S1.** Criteria of clinical sickness scores^1^

| **Clinical parameters** | **Degree** | **Score points** |
| --- | --- | --- |
| **Body weight loss** | Normal | 0 |
|  | Mil<5%) | 1 |
|  | 6-10% | 2 |
|  | 11-15% | 3 |
|  | 16-20% | 4 |
|  | >20% | 5 |
| **Appearance** | No piloerection | 0 |
|  | Piloerection | 1 |
| **Spontaneous behavior** | Alert | 0 |
|  | Slow moving | 1 |
|  | Lethargic | 2 |
|  | Immobile | 3 |
| **Eyes** | Normal | 0 |
|  | Squinted | 1 |
|  | Closed | 2 |
| **Provoked behavior** | Quickly moves away | 0 |
|  | Slow to move away | 1 |
|  | Do not respond | 2 |
| **Breathing** | Normal | 0 |
|  | Elevated | 1 |

<sup>1</sup> Adapted from Moreau et al. (Am. J. Trop. Med. Hyg., 2020) and Kumari et al. (Viruses, 2021).

**Table S2.** COVID-19 patient characteristics

| **Demographics** |  | **%** |
| --- | --- | --- |
| Number | 4 |  |
| Age (years) | 69.75±13.84 |  |
| Hospital day | 14.0±8.042 |  |
| Female | 1 | 25% |
| **Comorbidities** |  |  |
| Hypertension | 4 | 100% |
| Diabetes | 3 | 75% |
| Obesity | 1 | 25% |
| Lung disease | 0 | 0% |
| History of smoking | 2 | 50% |
| Heart disease | 2 | 50% |
| Kidney disease | 1 | 25% |
| Cancer | 0 | 0% |
| Autoimmune diseases | 0 | 0% |
| Immune deficiency | 0 | 0% |
| **Laboratory findings** |  |  |
| CRP (mg/dL)* | 9.968±4.828 |  |
| D-Dimers (µg/mL)** | 5.468±1.740 |  |
| LDH (U/L)# | 1,053.0±878.9 |  |
| Ferritin (ng/mL)& | 2,644±2,057 |  |
| Haemoglobin (g/dL) | 12.58±2.457 |  |
| Neutrophils (cell/mm<sup>3</sup>) | 7,475±4,518 |  |
| Lymphocytes (cell/mm<sup>3</sup>) | 1,850±1,396 |  |
| Platelets (count/mm<sup>3</sup>) | 160,000±71,643 |  |
| **Medications** |  |  |
| Antibiotics | 4 | 100% |
| Heparin | 4 | 100% |
| Antimalarial | 0 | 0% |
| Oseltamivir | 1 | 25% |
| Glucocorticoids | 2 | 50% |
| **Respiratory status** |  |  |
| Mechanical ventilation | 4 | 100% |
| Nasal-cannula oxygen | 4 | 100% |
| Room air | 0 |  |
| pO<sub>2</sub> | 59.98±17.50 |  |
| SatO<sub>2</sub> | 85.30±14.46 |  |
| **Outcome** |  |  |
| Deaths | 4 | 100% |

\*CRP: C-reactive protein (Normal value <0.5 mg/dL); \*\*D-dimers (NV <0.5 µg/mL); #LDH: lactic dehydrogenase (Normal range: 120-246 U/L); &Ferritin (NR: 10-291 ng/mL)

